# Netome: The Molecular Characterization of Neutrophil Extracellular Traps (NETs)

**DOI:** 10.1101/2020.05.18.102772

**Authors:** David Scieszka, Yi-Han Lin, Weizhong Li, Saibyasachi Choudhury, Yanbao Yu, Marcelo Freire

## Abstract

Neutrophils are the most abundant type of white blood cells in humans with biological roles relevant to inflammation and fighting infections. The release of neutrophil extracellular DNA aims to control invasion by bacteria, viruses, fungi, and tissue damage. Neutrophil Extracellular Traps (NETs) act as antimicrobial agents triggering immune signaling through the release of the nuclear content into the extracellular space. Although intense investigations have elucidated the pathways preceding NET formation, the exact molecular composition of released NETs has not been mapped. We aimed to decode the sequences of DNA and proteins from NETs. With emerging needs to understand neutrophil functions precisely, we open the field of NETOMIC studies through isolation of NETs in combination with omics approaches including shotgun genomics and proteomics. Our in vitro NET isolation methodology allowed for unprecedented replicability with induction in a sterile inflammation model system. Enrichment of mitochondrial DNA and telomere sequences are significantly expressed in NET genomes. This study revealed that the genomic sequence released in the extracellular milieu is not stochastically serving as a scaffold for a repertoire of proteins involved in neutrophil protective functions. Collectively, we established the gene and protein signatures exclusive to the extracellular NETs in comparison to undifferentiated and differentiated neutrophil states, further guiding future detection of specific regions needed for diagnostics and targeted therapies of NET related conditions.

## Introduction

As polymorphonuclear leukocytes of the phagocytic system, neutrophils are essential for early immune defense in sites of infection where they kill pathogens^1,2^. They are generated from myeloid precursors in the bone marrow prior to differentiating into several stages of maturation, including myeloblast, promyelocyte, myelocyte, metamyelocyte, bands, and polymorphonuclear cells. Neutrophils are the most abundant types of white blood cells in mammals - approximately 100 billion are produced in the human body every day. From the blood circulation, neutrophils are recruited by chemical and physical cues to the periphery and are able to quickly infiltrate tissues and interstitial areas to detect and eliminate threats. Neutrophils present key functions in the clearance of pathogens such as bacteria^3^, fungi^4^, viruses^5^, and parasites^6^. In addition to the early recruitment to infection sites and clearance mechanisms by phagocytosis, oxidative burst, and degranulation, another much less recognized means of cellular attack can develop. Web-like chromatin structures known as neutrophil extracellular traps (NETs) are produced by neutrophils and released from the intracellular regions for host protection and infection control^7,8^. The DNA backbone of NETs is attached to molecules, such as histones, calprotectin, and cathepsin G protease, which provide antimicrobial properties to eliminate invaders^9^. Neutrophil response and NET production require a fine balance: underactivity leads to increased invasion from pathogens, whereas over-activity is highly damaging to tissues. While NETs are favorable in the host defense against pathogens, secondary damage to tissue from sustained formation can lead to a cascade of inflammatory reactions, resulting in organ damage, cancer, tissue loss and thrombosis. When dysregulated, excessive NET release has been implicated in diseases, including lupus^10^, COPD^11^, type 2 diabetes^12^, chronic inflammation^13^, cystic fibrosis^14^, autoimmunity^15^, and cancers^16^, among others^17,18^.

Accumulating evidence supports the notion that expelling DNA to the extracellular environment is not generally considered a specialized immune function, it was thought that the release was not regulated, and that molecules involved were randomly placed in the extracellular environments. It is now accepted that this process is a fine-tuned and well-controlled intracellular process. Known factors guiding generation of NETs include neutrophil elastase (NE), peptidyl arginine deiminase type 4 (PADI4), and gasdermin D. The release is dependent on the type, concentration, and duration of stimulus presented^1,4,8,16,18,19^. Stimuli from microbes and chemicals act differently in activation, cell membrane rupture, and NET expulsion. In response to a more replicable substance, such as phorbol 12-myristate 13-acetate (PMA), common intracellular pathways are activated in neutrophils, including protein kinase C (PKC)-mediated pathways, and MAPK/ERK signaling which generates downstream reactive oxygen species (ROS) via NADPH oxidase and myeloperoxidase (MPO). In fact, MPO is believed to have several roles in NET formation pathways upon its release from azurophilic granules. Once activated, MPO can convert H_2_O_2_ into hypochlorous acid (HOCl), which is one of the most caustic chemicals used in NETs to terminate invading pathogens. MPO also utilizes ROS to mediate the activation of NE which translocates into the nucleus for initial histone degradation and chromatin unpacking. MPO further promotes chromatin unpacking by activating PADI4, which is responsible for the citrullination of histones. However, most of the scientific evidence was focused on the identification of intracellular pathways leading to NET release and not on dissecting the content expelled from neutrophils. There are multiple molecules released from neutrophils that are unknown, highlighting the need to decode the exact sequences of the human NETs.

Despite emerging interests in neutrophil phenotypes, their use in research experiments are limited due to the short lifespan of the cells, high sensitivity to handling and temperature, making it challenging to achieve replicability. To revisit these issues and determine NET composition, we devised a protocol that reliably produced NETs in a sterile inflammation setting. We leveraged a myeloid undifferentiated cell line (HL60 cell line) into our cell differentiations systems (dHL60). Through concentration-depended and time point PMA perturbation studies, we successfully transformed neutrophils into NET producing cells. Here, we aimed to identify the genomic sequence of NETs and to characterize their molecular contents (specific proteins, and metabolic markers for NET scaffold). We demonstrated through imaging, staining, sequencing, and bacterial clearance that the sequences were viable NET materials. The NET sequences were compared to undifferentiated neutrophils and differentiated neutrophils. Finally, we compared our data with a published dataset to understand the replicability of our model to other neutrophil models. Our study showed a replicable system to produce human NETs and identified the genomic and proteomic contents of the differentiated neutrophils. This first human sequence study determining the exact content material opens new avenues for elucidating NETs structures.

## Materials and Methods

### Experimental Design

To achieve our goal of creating an *in vitro* NET pipeline, promyeloblast human cell line HL-60 cells were selected due to their ability to differentiate into neutrophils. Kinetic studies were performed to validate the timeline of differentiation and different concentrations of PMA that were required to induce optimal NET release. These were validated through visual inspection, immunofluorescence, MitoSOX-red assays, NanoDrop, and Qubit.

### 1-HL-60 Cell Culture

Promyeloblast human cell line, HL-60, was acquired from ATCC. HL-60 cell lines were maintained in culture media prepared according to manufacturer’s guidelines. Briefly, the HL-60 culture media consisted of Iscove’s modified Dulbecco’s medium (IMDM, Gibco, cat. no. 12440-061) with 5% fetal bovine serum (FBS, heat inactivated, Gibco, cat. no. 26140-095) and 1X antibiotic (Penicillin + Streptomycin; Gibco, cat. no. 15-140-122). Note: Avoid using antimycotic in the recovery media as it affects the recovery and growth of HL-60 cell lines.

### 2-Differentiation of HL-60 to Neutrophil (dHL-60)

HL-60 cells in culture media were centrifuged at 275 x g for 10 minutes at room temperature (RT) and the culture media was aspirated. Cell pellets were resuspended in differentiation media (i.e culture media containing 1.5% dimethyl sulfoxide) to an initial cell seed count of 10E5 cells. Verification of a previous neutrophil differentiation timeline was performed^20^. To quantitatively determine the maximum number of differentiated cells before apoptosis, a growth curve was created using a 96-well plate measured daily for 5 days. Morphological changes were monitored by Giemsa staining through the comparison of HL-60 cells to differentiated HL-60 cells (dHL-60) i.e. neutrophils.

### 3-Neutrophil Extracellular Trap Production

Multiwell plates were used in kinetic studies to determine effective NET release from dHL-60 cells. PMA was used to induce NET release at concentrations ranging from 0.1 nM - 8,000 nM and at time points ranging from 10 minutes to 6 hours. Morphological changes were monitored through different microscopy techniques and DNA was quantified by NanoDrop spectrophotometer, Agilent High-sensitivity DNA chip using Bioanalyzer 2100 (Fig. 1E), and QuBit 2.0.

**Figure 1.**
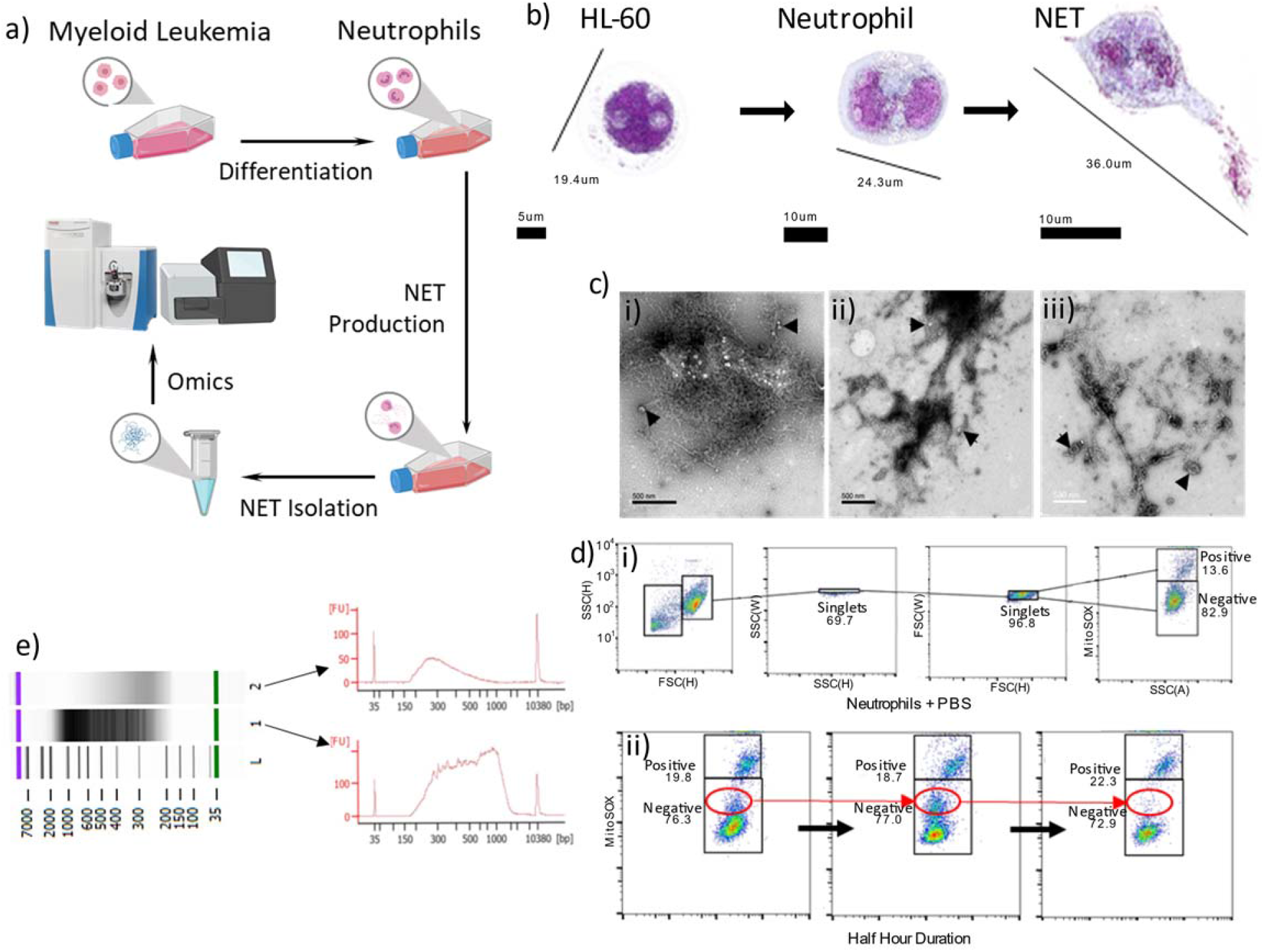
NET Production and Isolation. **(a)** Schematic of *in vitro* NET production. Cultured HL-60 cell lines incubated with DMSO treatment differentiate to neutrophils (dHL60 cells). After which, PMA stimulation leads to NET production, isolation, and “omics’’ analysis. **(b)** 3D holotomographic microscopy images, digitally stained based on RI (refractive index) confirms the differentiation of HL-60 cells to neutrophils (dHL-60) after 5 days in differentiation media and successful release of NETs after 4-hour incubation in 1,000 nM PMA. **(c. i-iii)** Electron microscope images of isolated DNA samples from **(i)** HL-60 cells, **(ii)** dHL-60 cells i.e. HL-60 and neutrophil DNA extracted via spin column and **(iii)** released NET extracted via centrifugation steps shows the presence of lipid bilayers in each sample (indicated by arrowheads). **(d)** ROS generation during NET release process was measured by MitoSOX assay that indicated Mitochondrial Superoxide formation over 5 hours of incubation with PMA. **(i)** Doublet discrimination gating strategy was used to ensure accurate MitoSOX-red quantification. **(ii)** Representative panel of flow cytometry analysis shows the generation of ROS from neutrophils on incubation with PMA for 2 hours. Red circle denotes the changes in the ROS generating neutrophil population over time. **(e)** DNA quantification by Agilent High-sensitive DNA chip verifies the composition of extracted NET samples. Lane 1 shows an isolated NET DNA sample. Lane 2 shows an isolated NET sample after incubation with DNase to digest all DNA contents. Arrows indicate the electropherogram of each sample in the gel image above. Lane L shows the DNA ladder marker.

### 4-DNA Isolation from HL-60, dHL-60, and NET

HL-60 and neutrophil (dHL-60) DNA was isolated via the Qiagen AllPrep DNA/RNA/Protein Mini Kit according to the manufacturer’s protocol^21^ and suspended in an elution buffer (EB) for storage at −20°C. DNA purity was quantified with NanoDrop spectrophotometer, QuBit 2.0, and Agilent High-sensitivity DNA chip.

We adopted a previous protocol for NET production from primary blood cells^22^. After 4 hours of incubation in PMA, culture media was aspirated to remove non-NET forming cells. Adherent, NET-releasing cells were resuspended in ice-cold PBS(-) to a final volume of 12 mL. The solution was centrifuged at 275 x g for 10 minutes to pellet cells. The NET-rich supernatant was transferred to an ultracentrifuge tube and the cell pellet was discarded. Ultracentrifugation was performed at 18,000 x g for 10 minutes at RT to pellet DNA and the PBS(-) supernatant was aspirated. NET DNA pellets were resuspended in EB for storage at −20°C.

### 5-MitoSOX staining and Flow Cytometry

The On-Chip Sort flow cytometer was used to quantify neutrophil ROS generation by staining with MitoSOX. Multiwell plates were used to compare HL-60 and dHL-60 cells in PBS(-) and in PMA over the course of 5 hours at 30-minute increments. In total, three experimental replicates were performed and analyzed.

### 6-Cell Staining and Microscopy

To monitor nuclear morphological changes, HL-60 and dHL-60 cells were stained with Eosin Y, Hematoxylin and Giemsa according to conventional methods and viewed under a histological microscope. Fluorescent staining was visualized using a Leica TCS SP5 II confocal microscope to qualify NET DNA release when compared to *E. coli* co-incubation. Both groups were permeabilized, stained with DAPI (1,000 nM), fixed with 4% PFA for 15 minutes, and checked for fluorescence. Additionally, nuclear morphology changes and NET release were visualized via 3D-Cell Explorer Nanolive microscope under fixed, and unfixed conditions. Comparative visualizations of isolated HL-60, dHL-60, and NET DNA was further performed using scanning electron microscopy.

### 7-Scanning Electron Microscopy

HL-60, neutrophil, and NET DNA samples were retrieved from −20° C and allowed to thaw on ice. Once thawed, samples were dehydrated in ethanol, embedded in epoxy resin, sectioned at 50– 60 nm on a Leica UCT ultramicrotome, and picked up on Formvar and carbon-coated copper grids. Sections were stained with 2% uranyl acetate for 5 min and Sato’s lead stain for 1 min.

Grids were viewed using a Tecnai G2 Spirit BioTWIN transmission electron microscope equipped with an Eagle 4k HS digital camera (FEI).

Formvar-carbon-coated copper grids (100 mesh, Electron Microscopy Sciences, Hatfield, PA) were placed on 20μl drops of each sample solution displayed on a Parafilm sheet. After allowing material to adhere to the grids for 10 minutes, grids were washed 3 times by rinsing through 200 μl drops of milli-Q water before being left for 1 min on 2% (wt/vol) uranyl acetate (Ladd Research Industries, Williston, VT). Excess solution was removed with Whatman 3M blotting paper, and grids were left to dry for a few minutes before viewing. Grids were examined using a JEOL JEM-1400Plus transmission electron microscope operating at 80 kV. Images were recorded using a Gatan OneView 4K digital camera.

### 8-Proteomics

Isolation of HL-60 and dHL-60 protein was conducted via the Qiagen AllPrep DNA/RNA/Protein Mini Kit and conducted according to the manufacturer’s instructions^21^. Attempts using the AllPrep to isolate NET protein from total NET samples were unsuccessful and the total NET isolation protocol was used for proteomics analysis.

### 9-NET protein preparation for proteomics

After thaw from −80°C, NET solutions were added with protease inhibitors and 1% Benzonase Nuclease (Sigma) and incubated at 37°C for 20 min to remove nucleic acids. Protein samples were then processed using the Suspension Trapping (STrap) approach as described previously^23^ to generate tryptic peptides for liquid chromatography-tandem mass spectrometry (LC-MS/MS) analysis. To specifically identify nucleic acid-bound proteins, one set of NET samples were processed similarly but excluded the Benzonase treatment.

In total, LC-MS/MS was performed on 6 protein samples (three NET with Benzonase treatment and three NET without Benzonase treatment) following a protocol described previously^24^. In brief, the desalted samples were first resuspended into 20 μl 0.1% formic acid in water and then loaded onto a trap column (2 cm × 300 μm, PepMap C18, Thermo Scientific) and an analytical column (19 cm × 75 μm, 3.0 μm; ReproSil-Pur C18-AQ media) coupled to an Ultimate 3000 nano-LC and Q-Exactive mass spectrometer system (Thermo Scientific). Protein identification and quantitation were performed using Proteome Discoverer (version 2.2) and MaxQuant-Perseus software suite. The UniProt human database (20,413 sequences, reviewed only; version 2018_12) was used for protein search. Only the peptide and protein identifications with false discovery rate (FDR) of 1% or less were accepted in the final data set. More details of proteomic procedures can be found in the recent publication^24^. The mass spectrometry proteomics data have been deposited to the ProteomeXchange Consortium via the PRIDE partner repository with the dataset identifier PXD016143.

### 10-Genomic Sequencing

Illumina’s NexteraXT library prep kit was used for HL-60, dHL-60, and NET samples and sequenced on the NovaSeq 6000 platform using an S2 flow cell 2X150bp. 400 pM of each sample pool was loaded and 1% PhiX was spiked in each lane^25^. Cluster density was 2961K/mm2 with 80% PF. 1429.42Gb and 4.5B PE Reads were generated. Coverage for the HL-60, dHL-60, and NET samples were 36X, 45X, and 47X respectively. The raw genomic sequences are available at NCBI Sequence Read Archive (SRA) under BioProject under accession PRJNA587717 at https://www.ncbi.nlm.nih.gov/bioproject/PRJNA587717.

The resultant fastq files were input into the CLC Workbench (v11), trimmed by 15 nt on the 3’ to remove primers, trimmed for q-scores < 25, mapped to the human genome (hg38, CLC v12), and checked for normal human GC content. To determine genomic enrichment, a sliding window analysis was conducted for every 20, 100, 500, and 5,000 nt. Regions of interest (ROIs) were selected based on known protein association^9^, proteomics results, and nonselective general analysis. Sliding window files of aforementioned sizes are available through the corresponding author. Additionally, unmappable read files are also available and were not used in our analysis.

#### Normalization Steps for Genomics Sliding Window

Normalization of sliding window coverage was performed by dividing coverage per sliding window by complete coverage per sample per chromosome.

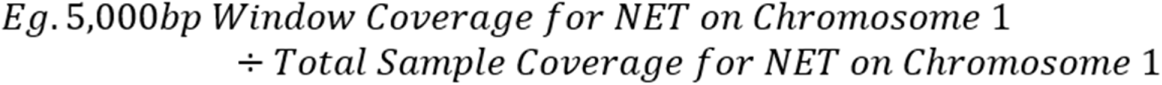

Normalization of mitochondrial DNA was performed by dividing coverage per sliding window by complete coverage per sample for the entire genome.

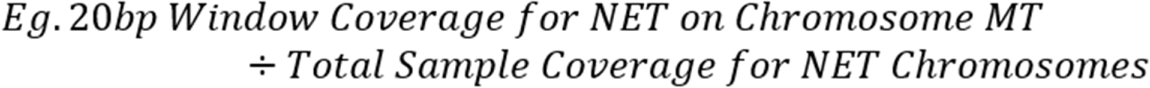

#### Known Protein Association

Proteins known to be localized within NETs were tested for genomic enrichments or depletions based on normalized 20 nt sliding windows^9^. These proteins of interest (POIs) were input into Ensembl biomart (Release 96) and their genomic start/end locations were determined.

#### Proteomic Protein and RNA Expression Selection

Proteomics analysis was used to retroactively determine ROIs. The genomic regions associated with the protein results were input into Ensembl biomart and their genomic start/end locations were determined. Enrichment/depletion analysis was performed with 20 nt normalized sliding windows.

#### Nonselective General Analysis used for RNA Expression Overlap

Using the 5000 nt normalized window analysis, regions were selected based on 1.5-fold-change for each sample compared to the others simultaneously or compared to the average. The annotated list used for exonic analysis was acquired from Ensembl biomart and used in conjunction with the normalized 5,000 nt windows. Overlapping expression analysis was performed by downloading the neutrophil gene expression data table (http://collinslab.ucdavis.edu/neutrophilgeneexpression/), and comparing the gene list generated from our nonselective general analysis. The initial range of genomic coverage spanned several orders of magnitude and a percentage of the total became more representative for visualization. Namely, each sample was divided by the average of all three samples.

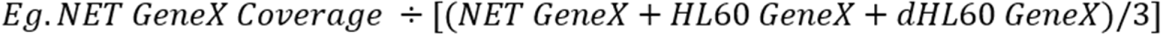

It should be noted that this normalization was simply for aesthetic reasons (Fig. 3) and was not performed during our initial genome-wide scan. All circos plots were generated using the ShinyCircos software (http://shinycircos.ncpgr.cn/).

Neutrophil elastase fluorescence and growth curves were monitored using the Celigo S cell imaging cytometer. Briefly, coculture experiments with *Fusobacterium* were performed in 6-well plates at a 1:10 ratio of neutrophils to bacteria, respectively. Standard curves were generated according to manufacturer instructions. Celigo internal cell counting software was utilized to determine cell growth each day over 5 days to determine cell division and terminal differentiation in 6-well plates.

Statistical significance was calculated with a nonparametric Mann-Whitney test and Prism software (Graphpad Software Inc.). Statistical significance for proteomic dataset was calculated with ANOVA. Genomic statistical significance for NETs, telomeric frequency, and mitochondrial enrichments were calculated using ANOVA. Statistical significance for correlations was calculated by Spearman rank test with P values and r values noted on the respective graphs. *P<0.05, **P<0.01, ***P<0.001, and ****P<0.0001 were considered significant and are referred to as such in the text.

## Results

### Neutrophil Differentiation and NET Release Model

To develop a neutrophil differentiation protocol and produce NETs *in vitro*, we first established a differentiation system that allowed cells to be viable with synchronized NET production. In mimicking human neutrophil differentiation, cells need to be primed prior to a specific response such as oxidative burst, phagocytosis or NET^26^. Extensive literature related to HL60 cells have made this an attractive model for studies of differentiation. Because neutrophil differentiation protocols lead to increased cell death, we established a threshold of differentiation that would keep viability at 80%. HL60 cells grow in suspension culture with a doubling time that can vary from 20-45 hrs. Morphologically, the cells present with rounded nuclei and basophilic cytoplasm with azurophilic granules (Figs. 1B, S1A). The majority of the cells carry a variety of histochemical markers characteristic of myeloid lineage, most notably myeloperoxidase and acid phosphatase. We evaluated to confirm cell morphology histochemically through trypan blue, Giemsa, and by flow cytometry analysis (CD11b). We initiated testing cell differentiation by culturing undifferentiated HL60 with culture medium enriched for DMSO (0.1-10%). As compared to steady state, neutrophil differentiation lead to cytoplasm enlargement, loss of primary granules, emergence of secondary granules, nuclear condensation and segmentation. We then established neutrophil differentiation at the viability threshold, with morphological characteristics and surface marker validation at 5 days under 1.5% DMSO (Figs. 1B, S1A). As expected, cells were terminally differentiated and presented phenotypic characteristics of molecular neutrophils.

One of the key characteristics to discriminate neutrophil differentiation to monocyte lineage from the progenitor cells is their nuclei shape and production of classic neutrophil enzymes such as elastase, myeloperoxidase, and ROS formation. We therefore further confirmed that our cells followed the fate of mature neutrophils rather than macrophages. Once HL-60 cells were differentiated into mature neutrophils, we performed kinetic studies to investigate NET release. Our pilot experiments with live imaging cytometry assayed the optimal concentrations of PMA stimulation (1-10,000nM). Previous data indicated that higher and lower concentrations under longer periods of time were able to induce NETs^27^. In our system we achieve synchronized NET formation by applying PMA (1,000nM). In the same experiment we investigated the kinetics to understand the ideal time for NET release (0-6 hours). Our morphological analysis demonstrated that at 4 hours cells were viable with morphological changes leading to spike NET formation. It is known that prior to NET release, mitochondrial superoxide is formed along with reactive oxygen species. We evaluated our cells by either morphological analysis or MitoSOX-red staining of mitochondrial superoxide (Figs. 1B, D, S1A, B). At 2.5 hours, our results showed a peak of ROS formation leading to increased staining of MitoSOX (Figs. 1D, S1B). Comparatively, neutrophils in PMA were 75.23% positive for MitoSOX versus 14.98% in PBS while HL60 cells in PMA showed 23.05% positive versus 10.89% in PBS. These early events preceded NET formation and were replicable through flow cytometry monitoring. Quantitatively, stimulation increased the number of oxygen species which allowed the separation of cells into three groups: MitoSOX^lo^, MitoSOX^Mid^, MitoSOX^high^, with intermediary subsets of MitoSOX^high^ representing the producers of high amounts of NETs.

The phenotypic identity of neutrophils was further confirmed through viability staining (SYTOX and DAPI). While the expression levels of neutrophil proteins were not affected by the high PMA 1,000 nM concentration, the viability decreased to 50% at 5 hours and 30% at 6 hours. It is possible that to produce NETs from leukemia cell derived cells, the genetic profiling is different when compared to primary cells which have low viability in *ex vivo* settings. Upon selection criteria determination for NET induction and release (4 hours and 1,000 nM) of differentiated cells, we proceeded to isolate NETs by adapting a protocol previously described^28^. We confirmed that a minimum neutrophil count of 1.0E6 was needed for measurable DNA isolation. We next confirmed the expression of DNA release by nucleotide staining, gel analysis, spectrophotometry and scanning electron microscopy (SEM). We controlled our assays by comparing the NET isolated groups with the undifferentiated cells and differentiated cells. Clearly, the NET groups present morphology and readings to DNA. To ensure our NET materials presented with isolated DNA from the extracellular sites and not from the cells, we visualized the isolated materials through SEM. The only group demonstrating thread-like structures and weblike structures was the NET group from the isolation layer – acquired from NETs adherent to the culture flask after PBS wash – while other phase layers of the high centrifugation protocol failed to demonstrate these patterns. To confirm the molecular content, DNase was applied to the isolated samples and compared to DNase free isolates (Fig. 1E). When we applied DNase to the material released, we showed that quantitatively and through imaging that the structures and DNA diminished, further confirming the nucleotide nature of NETs. The presence of lipid bilayers in SEM images of NETs could also be vesicles, which are commonly known to be released by neutrophils (Fig. 1C). Collectively we defined a method to differentiate neutrophils, and to produce and isolate NETs. Following our validation protocols, we proceeded to extract DNA/RNA/protein from our three groups for proteomics and genomics sequencing.

### NET Proteome Analyses

The analysis of proteomics data revealed significant differences on the NET groups compared to undifferentiated and differentiated neutrophils in expression levels of proteins known to be associated with neutrophil function and NETs. To ensure the protein content was from NET and not from cells, our system allowed us to investigate proteins that were exclusive to NET groups and not the cell groups. Principal component analysis showed that NETs presented unique clusters when compared to cell proteins (Fig. S4). Out of initial 2,403 proteins quantified in NETs, 316 proteins were enriched when compared to neutrophils and undifferentiated cells (HL60; Fig. S5). Although the isolated NETs presented with unique and overlapping sequencings, this group was the only one to demonstrate enrichments for classic pathways for NET formation, confirming our DNA and cytometry findings. In line with the specific protein signatures, differentiated and undifferentiated cells showed reduced numbers of NET markers and presented the highest overlapping rates with pathways related to extracellular exosomes, mitochondrial metabolism, mitochondrial nucleoid, and extracellular matrices (Fig. S5).

We next assessed isolated samples from NET groups and cells were processed for shotgun proteomic analyses to characterize the protein identity and function. One set of samples were directly processed using the established method to generate tryptic peptides for LC-MS/MS analysis^24^, while the other set of samples were treated with Benzonase to degrade all DNA and RNA contents prior to tryptic digestion. Because Benzonase is able to cleave the condensed nucleic acid component in collected NET, we monitored its activity to subsequently release proteins that are tightly bound to the nucleic acids, thus enhancing proteomic mapping^29^. Without Benzonase treatment, the abundance of protein levels was moderate with 1,711 proteins identified. Whereas after Benzonase treatment an increased abundance of proteins was found, with identification of 2,364 proteins (Fig. 2A). The proteome size spans higher than five orders of magnitude, and in addition to novel protein findings, the protein functions previously associated with NET in comparison to our study findings is listed (Fig. 2B)^9^, further validating the enrichment of NETs in our culture and isolation system.

**Figure 2.**
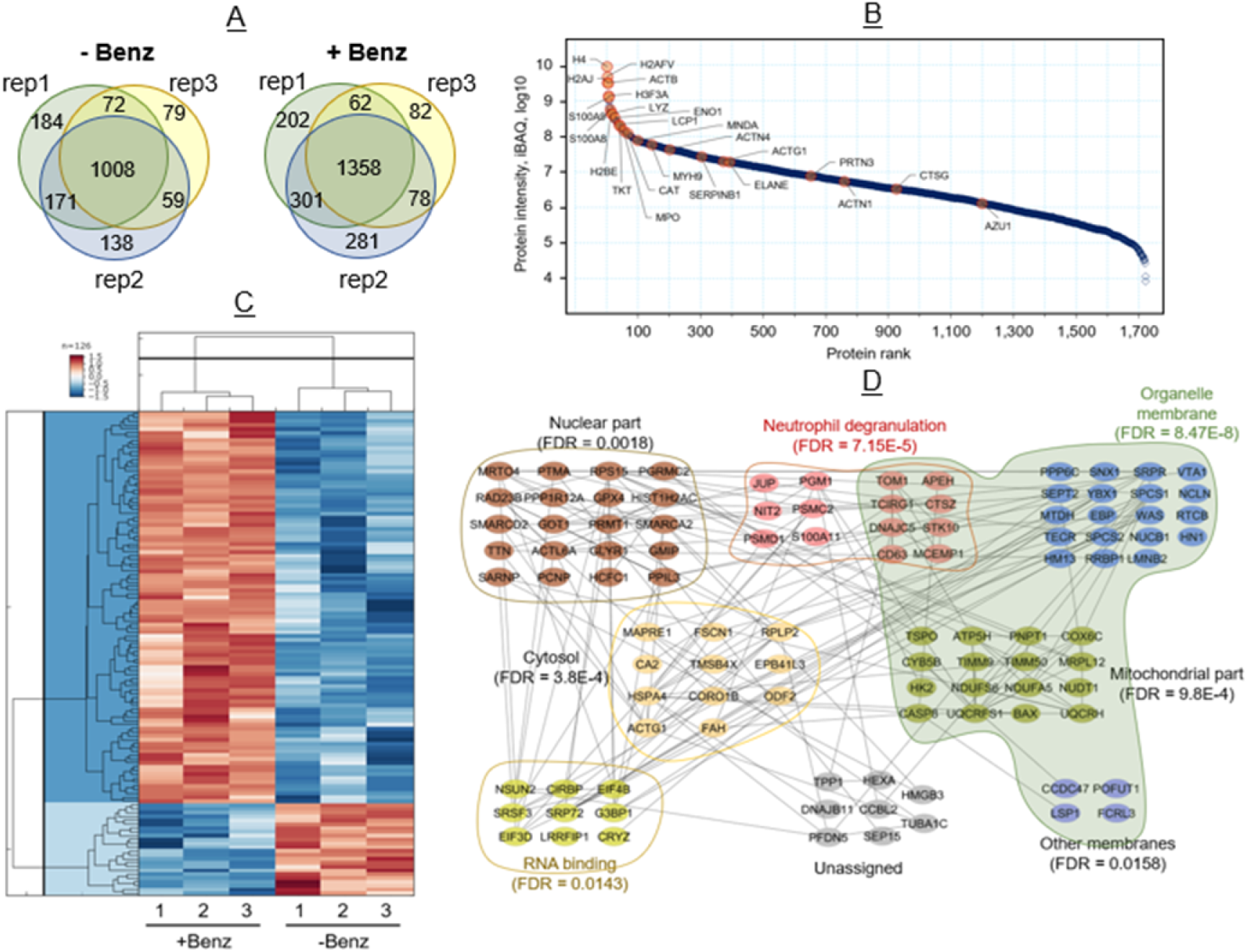
NET Proteome. **(A)** Proteome analysis of NET from three representative samples (i.e. rep1, rep2, and rep3) identified a total of 2,364 proteins after Benzonase treatment and 1,711 proteins in untreated samples. Common proteins found among three representative samples in Benzonase treated and untreated NET is 1,358 and 1,008, respectively. **(B)** Dynamic range of the NET proteome. Data showing (1,722 proteins) here is from Benzonase-treated NET. Median values of the three replicate experiments were used for the plot. Previously reported proteins associated with NET by Urban *et. al*. are denoted by orange dots, most of which ranked among the 100 most abundant proteins found in our NET samples. (Median value of three experimental replicates are plotted here). **(C)** Hierarchical clustering of the 126 significant proteins (fold change ≥ 2 or ≤ −2; Permutation FDR 0.05) between the two groups. Z-scored LFQ intensities were color coded as indicated in the scale bar. **(D)** STRING protein network and Gene Ontology analysis of 101(out of 126) significantly enriched proteins in NETs after Benzonase treatment were analyzed using embedded STRING app in CytoScape software (version 3.7.2). The confidence score cutoff was set to 0.4. Representative enriched gene ontology (GO) terms (e.g., biological process, molecular function and cellular compartment) and corresponding FDR values were depicted in the network.

Functionally, the 100 most abundant proteins identified in our proteome analysis included histone proteins H4, H2AFV, H2AJ, and H3F3A, which are known to be the core components of nucleosome and play a central role in cell functions including transcription regulation, chromosome stability, DNA repair and DNA replication. Consistently, a classic NET enzyme, Myeloperoxidase (MPO), which was evaluated by flow cytometry in our culture experiments was also found in high abundance. This was also evident for the calcium and zinc-binding proteins known to be released by blood neutrophils in humans after migrating to the site of inflammation, calprotectin (S100A8 and S100A9), which plays an important role in the regulation of inflammation and immune response^30^.

The Label Free Quantitation (LFQ) intensities correlate well within the same treatment groups (Fig. S6), showing consistent reproducibility of NET proteome profile. Overall, 2,280 proteins were common within the NET groups, and 126 proteins have more than two-fold difference between the two groups by statistical analysis (Permutation FDR 0.05, Fig. 2C). Histone proteins, cathepsin G, S100A8, S100A9, and azurocidin have more than two-fold increase in the Benzonase-treated group, indicating their tight association with the condensed nucleic acid component in NET. Upon functional analysis of the 101 proteins with at least two-fold increase in the Benzonase-treated group, we found that while a fraction of the proteins is involved in leukocyte activation and neutrophil degranulation (FDR= 7.15E-5), another group of proteins related to RNA binding (FDR= 0.0143) was also enriched (Fig. 2D). We attempted to extract RNA from NETs, but were not able to isolate detectable levels. However, the proteomic findings indicate that the condensed nucleic acid component from NET may not only be the chromosome scaffold, but also includes cellular RNAs.

Given the abundance of novel proteins, we sought out alternative data organization methods for pathway enrichment analyses to test against our subsequent genomic sequencing. As such, we organized the proteomic score by peptide spectrum matches (PSM) and input the resultant protein hierarchy into the panther gene ontology pipeline (http://www.pantherdb.org/), which demonstrated pathways associated with positive regulation of neutrophil degranulation, cytosol, mitochondrial proteins and RNA binding molecules. In addition to the abundance of mitochondrial proteins, NETs contained a statistically significant increase in the telomere sequence TTAGGG from our genomics sequencing (Fig. 4B). To our knowledge, this is the first evidence that neutrophils could be increasing the amount of DNA to be released in the NET process *via* the extension of telomeres through the internal RNA template of human telomerase reverse transcriptase.

### NET Genome Analysis

#### Sliding Window Analysis for Non-Targeted Genome-Wide Scan

To interrogate the genomic sequence of the scaffold DNA released by neutrophils, we utilized Illumina’s NexteraXT library prep kit and sequenced NET genome on the NovaSeq 6000 platform. Coverage for isolated Nets were compared to undifferentiated cells and differentiated cells (36X, 45X, and 47X respectively). The raw genomic sequences were submitted to NCBI Sequence Read Archive (SRA) under BioProject under accession PRJNA587717 at https://www.ncbi.nlm.nih.gov/bioproject/PRJNA587717.

The resultant fastq files were input into the CLC Workbench (v11) software, trimmed by 15 nt on the 3’ end to remove primers, trimmed for q-scores < 25, mapped to the human genome (hg38, CLC v12), checked for normal human GC content, and comparatively scanned using a sliding window analysis^31^ which was normalized to each sample’s chromosome. To determine genomic enrichment, a sliding window analysis was conducted for every 20, 100, 500, and 5,000 nt, with the 500 nt represented for mitochondrial DNA (Fig. 4D) and the 5,000 nt results represented for whole genome analysis (Figs. 3, S2). Regions of interest (ROIs) were selected based on known functions^9^, proteomics results, and nonselective general analysis. Unmappable read files were not used in our analysis. From the sliding window approach, we were able to reveal regions of enrichment and depletion within each sample.

**Figure 3.**
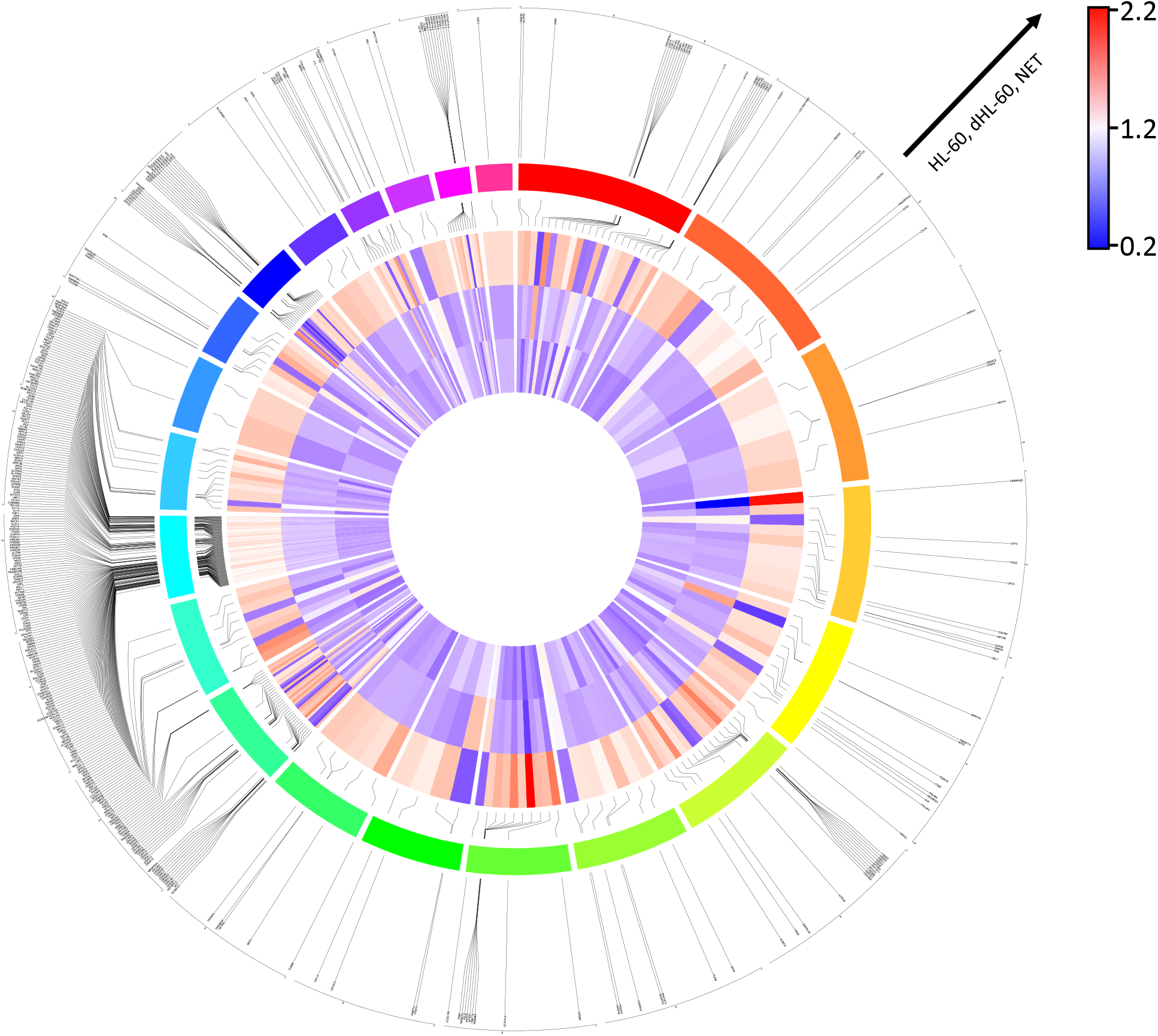
NET DNA Enrichment/Depletion Throughout Human Genome. A 1.5-fold cutoff enrichment screen was used to determine regions of enrichment/depletion. Using annotated gene coding regions, a comparison to published expression data was used to determine overlap. The resultant circos plot for chromosomes 1-22 is shown. From innermost to outermost tracks: heatmap in order of HL-60, neutrophil (dHL-60), and NET enrichments; linkers from heatmap to color coded chromosomes in order from red (chromosome 1) clockwise to pink (chromosome 22); and linkers to gene names.

After the genome-wide scan was conducted, comparative analyses were made for each possible grouping of samples using a 1.5-fold enrichment/depletion cutoff. From this, each sample grouping revealed many intronic and exomic regions of interest. Most notably, there were 25,488 regions in the NET when compared to the HL-60 and neutrophil sample groups (Fig. S3F), whereas HL-60 showed this in 6,147 regions (Fig. S3D) and neutrophil showed this in 5,351 regions (Fig. S3E) Overall, there were 538,355 regions within this analysis that showed no consistent enrichment or depletion when compared to any possible grouping (Fig S3G). We retrieved the genes associated with the 25,488 regions and represented them in figure S2. These data could suggest that NET DNA scaffold is being released in a time-dependent manner due to the regions of NET enrichment and depletion being 4-fold higher than neutrophil or HL-60, with HL-60 and neutrophil regions being relatively similar to each other.

We reasoned whether DNA was being released in an ordered fashion, i.e. DNA not deemed essential for immediate survival or NET release was being preferentially ejected first. To rationalize the sequence of NETs with neutrophil function, we looked into known RNA-seq datasets related to similar cell lineage. In the case of NET release, the expression of survival genes would necessarily require them to be inside of the neutrophil as other DNA is being ejected. To ensure our sequences align, we probed previously published data HL-60 undifferentiated and differentiated RNA-seq data deposited at the NCBI Gene Expression Omnibus (GEO) database (Accession: GSE103706) along with previously published data for primary human neutrophils from the Wright et al [GEO: GSE40548], Jiang et al [GEO: GSE66895]. This comprehensive list was compared to our NET enrichment/depletion list. Our findings demonstrated that out of a list of 37,470 RNAs, 400 overlapping regions were determined (Fig. 3), indicating enrichments and depletions of NET genes present in the extruded materials.

### Sliding Window Analysis for Targeted Candidate Gene Regions

We next sought to ascertain what DNA sequences were enriched in NET samples when compared to the neutrophil groups. We performed an additional targeted candidate analysis of genes related to the known NET release pathway(s). Previous studies of genes known to be implicated in NET release showed statistically significant DNA depletion in our NET samples, such as PADI4 and MPO (p < 0.05, Fig. 4A). We investigated genomic telomere enrichments using targeted window analysis (TTAGGGTTAGGG). Telomere counts in NET sequences showed statistically significant increase over undifferentiated and differentiated neutrophil telomere counts, while HL-60 and neutrophil telomere counts were not significantly different from each other (p < 0.05, Fig. 4B). We further identified that mitochondrial sequences showed significant enrichment in NETS when compared to HL60 and neutrophil counts, whereas HL-60 and neutrophil mitochondrial enrichments were not significantly different from each other (p < 0.05, Fig. 4C). After a positional mapping comparison, the mitochondrial regions of specific NET enrichment were determined to be the NADH dehydrogenase subunit components ND1, ND2, ND4, and ND5 (Fig 4D). These data suggest that PMA-induced NET release is comprised of mitochondrial and nuclear DNA and that neutrophils could be upregulating telomerase in order to maximize the amount of DNA being released into the extracellular space. Taken together, these could suggest an ordered release mechanism that preferentially ejects DNA not deemed necessary for the NET release process before complete DNA ejection takes place.

**Figure 4.**
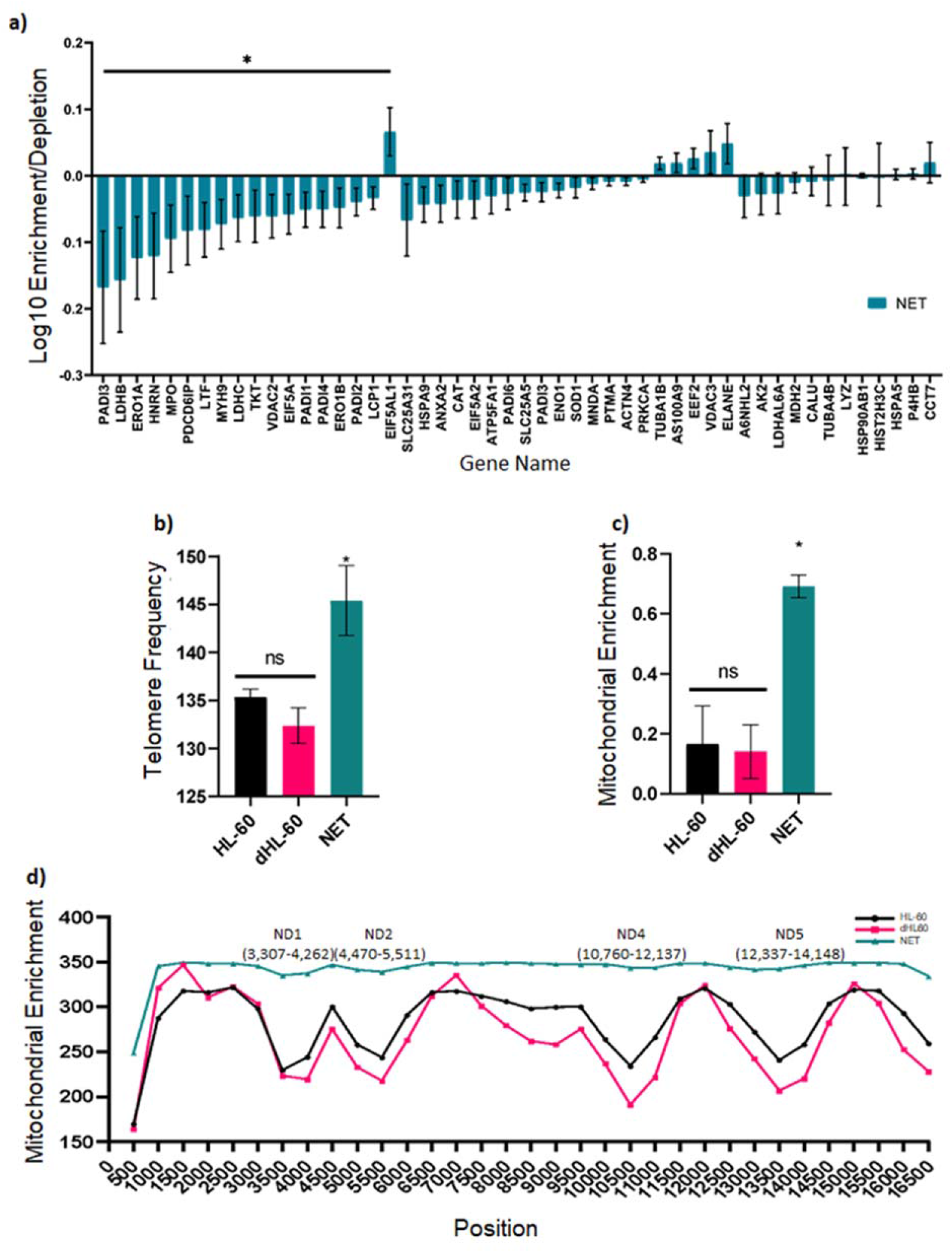
NET Genomic Enrichment Regions. All comparisons are made between HL-60, neutrophils (dHL-60), and NET samples. **(a)** *A priori* gene comparison based on proteomics results, previously reported proteins by *Urban et al*., and known biochemical pathways associated with NET formation. (p-value: * <0.05) **(b)** Comparison of telomere counts (TTAGGGTTAGGG) across all chromosomes for each sample (ANOVA, p < 0.05). **(c)** Normalized mitochondrial enrichment quantification from two sequencing runs (n = 2 per sample, ANOVA, p < 0.05). **(d)** Mitochondrial enrichment by position number (sliding window size = 500nts).

To validate that our NETs were functional and presenting antimicrobial properties, we incubated with a pathogenic strain of gram-negative microbe (*Fusobacterium*) and investigated the microbial viability and the rate of NETs *via* elastase. As observed in *ex vivo* and *in vivo* settings^32^, NETs were detectable earlier than the 4 hours required by PMA. Through microscopy, we monitored NET formation timepoints and observed an increase of NET formation through a survey of markers for NET quantification. Expression of neutrophil elastase was increased with *F. polymorphum* over neutrophils in PMA, while neutrophils in PMA had increased expression over our negative control (neutrophils in PBS), with the other strains of *F. nucleatum* and *F. animalis* having no difference when compared to neutrophils in PMA (Fig. S1C). Thus, NET formed *ex vivo*, augmented with specific bacterial strains facilitate inflammation activation and bacterial clearance over 4 hours.

## Discussion

An excessive and persistent neutrophil response is involved in the pathology of an array of human conditions, and NETs have consequently become an attractive therapeutic target. Yet, therapeutic strategies looking into improving pathogenic inflammation have failed to fully consider the potential role of DNA sequences released by these cells. Mature neutrophil differentiation is often associated with increased cell death and NET release. We set out to determine the protocol *in vitro* that mostly produced NETs with replicability. We compared most commonly used differentiation agents including DMSO, PMA, LPS, formaldehyde (Fig. S1A), selected DMSO for differentiation, and tested the NET release in dose-response and time-dependence (0-6 hours) through microscopy and by early (MytoSOX) and late differentiation markers (CD11b). By monitoring their ability to differentiate from HL60 to dHL60 we were able to characterize their morphology precisely along with decrease in cell size, increased nuclear pyknosis, and segmentation along with marker expression (Figs. 1, S1A). This *in vitro* differentiation made the HL-60 cells ready for NET formation at 4 hours in 1,000 nM PMA. Throughout the course of our NET studies, we also had the opportunity to visualize an unparalleled amount of neutrophil NET formations in response to sterile inflammation and co-culturing experiments with different strains of bacteria (Fig. S1C). From these, several trends emerged, such as the seemingly directed NET release towards bacterial colonies when present, and NET release perpendicular to culture media flow in the presence of PMA.

To further assess the degree of differentiation of the cells, we assessed nuclear size and morphology through live imaging in a holotomography model. The cells were further isolated and processed for proteomics. Although further studies are needed to confirm the molecular composition of sterile PMA-induced NET release, our initial proteomic results confirm the existence of known NET-associated proteins. The origin of NET DNA remains controversial, with studies reporting nuclear, mitochondrial, or both depending on the cell type and stimulus^33–36^. Here, the existence of mitochondrial enrichment as well as nuclear DNA adds to the body of evidence showing PMA-induced NET release has origins in the nucleus as well as mitochondria. Protein comparisons of NETs to cells in both proteomics (Figs. S4, S5) and genomics (Fig. S3) demonstrated that mitochondrial metabolism was highly enriched.

The proteomic studies, specifically, demonstrated that NETs are associated with a significant amount of proteins. The greater amount of proteins identified from Benzonase-treated NETs versus the untreated ones (2,364 vs 1,711) indicates that many proteins interact tightly with the nucleic acids while being released from the neutrophils. Proteins significantly increased after Benzonase treatment are enriched in those involved in neutrophil degranulation, nucleus and mitochondrial proteins, and membrane proteins (Fig. 2D). Although not investigated in this paper, it is interesting to ponder whether Cdc42 and GPCRs could be involved in a directional NET release depending on environmental cues such as shear forces or chemoattractants. NET formation pathways are still under investigation, and the complexity of understanding these processes relate to the fact that neutrophils are short-lived cells and not amenable to gene editing. Alternatively, strategies that seek to promote neutrophil survival and to modulate NET release to reduce inflammation, but simultaneously activate a more robust innate immune response, will leverage the understanding of NET sequences in the extracellular niche. Emerging pathways will be found from replicable approaches such as those utilized in this report. Thus, in addition to revealing the number and type of proteins attached to the NETs we have also categorized them according to common neutrophil functions.

Our initial study of the molecular content within NETs has opened many questions, including cellular processes that increase the amount of DNA being released through telomeric expansion as well as region-specific NET release for prolonging cellular responsiveness. With the understanding that there are at least two types of NET release - vital and suicidal^37^ - we reasoned that the neutrophils could have secondary internal recognition pathways which could preferentially sequester genomic regions for maximal longevity. PMA typically involves suicidal NET release. However, the sequestration of genomic scaffolding could be time dependent, meaning that certain regions could be consistently released before others. Our findings of specific NET release-associated genomic scaffold depletions (Fig. 4) could imply this to be true. Further exploration utilizing super resolution microscopy and DNA painting could elucidate these mechanisms with greater resolution. Additionally, the finding of significant telomere enrichment within the NET samples was surprising, but an interesting and unexplored avenue. In conjunction with the DNA sequestration rationale, we reasoned that neutrophils could be increasing the internal stockpile of DNA before ejection toward invading pathogens. From the perspective of biological load, this would allow for maximum efficiency of pathogen binding and immobilization at a lower cost relative to the complete elimination of a second neutrophil undergoing NET release. Additional studies are needed to survey the specificity and diversity of NETs in the context of their niches and cell types, while taking into consideration differentiation states, health, and disease.

Finally, we wanted to compare our results obtained with the dHL60 cells induced NET formation to a database of neutrophils^38^. This validation showed that our protocol had optimal differentiation by using PMA. When analyzing the global gene expression in myeloid cell lines a total of 400 genes were similar to RNA-seq data from undifferentiated and differentiated cells. To account for measurement noise, we repeated our sequencing experiment in three independent samples. For comparison, we collated raw datasets for our cell lines with the dataset from previously published studies and analyzed these data sets along-side our own using identical parameters and normalization strategies (Fig. 3). The availability of this *in vitro* model holds promise for studies into disease states, as well as probiotic discovery through bacterial factors that modulate NET release, as seen by the differing amounts of NET release during different bacterial cocultures.

Together, our observations reveal that the genomic detection provided the scaffold information by which NETs release functional proteins to the extracellular milieu. PMA was sufficient to produce replicable amounts of NET in a dose response manner, impacting the results of our proteomics and genomic analysis. Our NET database will be a valuable tool to identify similarities and differences for gene and protein expression between primary and cell lines and improve our understanding of NET biology and pathology.

**Table 1.** List of Reagents and Platforms.

| *Reagent/Item Description* | *Reference number* |
| --- | --- |
| HL-60 cell line | ATCC, cat. no. CCL-240 |
| IMDM | Gibco, cat. no. 12440061 |
| FBS | ATCC, cat. no. 30-2020 |
| Penicillin + Streptomycin | Gibco, cat. no. 15140122 |
| Cell Imaging Cytometer | Nexcelom, CeligoS, cat. no. 200-BFFL-S |
| Agilent Bioanalyzer 2100 | Agilent, model G2939A |
| QuBit 2.0 | Invitrogen, cat. no. Q32866 |
| Qiagen AllPrep DNA/RNA/Protein Minikit | Qiagen, cat. no. 80004 |
| Ultracentrifuge tube | cat. no. 149569C |
| On-Chip Sort Flow Cytometer-2D Chip-Z1001 | cat. no. 1002004 |
| MitoSOX™ Red Mitochondrial Superoxide Indicator | Invitrogen, cat. no. M36008, lot 2015529 |
| Eosin Y, 0.25% (w/v) in 57% Alcohol | Ricca Chemical, cat. no. 284516 |
| Hematoxylin Stain Solution, Gill 1 formulation, regular strength | Ricca Chemical, cat. no. 353516 |
| Giemsa Stain Solution | LabChem, Inc., cat. no. LC148407 |
| Confocal Microscope | Leica TCS SP5 II |
| Cytofix/Cytoperm Fixation and Permeabilization solution | BD Biosciences, cat. no. 55472251-2090KZ |
| DAPI | Invitrogen, cat. no. D1306 |
| Cytoseal-60 mounting media | Thermo Scientific, cat. no. 83104 |
| Tomographic Microscope | Nanolive 3D Cell Explorer |
| Scanning Electron Microscope | Leica UCT Ultramicrotome |
| Grid Viewing Microscope | Tecnai G2 Spirit BioTWIN transmission electron microscope equipped with an Eagle 4k HS digital camera (FEI) |
| Formvar-carbon-coated copper grids | 100 mesh, Electron Microscopy Sciences, Hatfield, PA) uranyl acetate (Ladd Research Industries, Williston, VT |
| Grid Examination | EOL JEM-1400Plus transmission electron microscope operating at 80 kV |
| Grid Recording Camera | Gatan OneView 4K |
| NexteraXT library prep kit | Illumina, cat. no. FC-131-1096 |
| Sequencing Platform | Illumina, NovaSeq 6000 |
| Flow Cell | S2 2X150bp |
| DNA Mapping Software | CLC Workbench v11, v12 |
| Neutrophil Elastase ELISA | Abcam, cat. no. ab204730 |

## Supporting information

u_46267_0_supp_1031906_q7mzdz_version2

## General

We would like to thank the J. Craig Venter Institute Sequencing Core for their tireless efforts to provide clean sequencing data and storage support. We would also like to thank Timothy Meerloo, Director of the UCSD Electron Microscopy Core for assistance in sample preparation and training.

## Author contributions

M.F. conceptualized, designed the study. D.S. performed experiments and partial bioinformatics analysis. W.L. performed genomic data analysis. Y.Y., YH.L. performed proteomics experiments and performed data analysis.

All authors wrote the manuscript and D.S and M.F edited for clarity.

## Data and materials availability

The mass spectrometry proteomics data have been deposited to the ProteomeXchange Consortium via the PRIDE partner repository with the dataset identifier PXD016143. The raw genomic sequences are available from NCBI Sequence Read Archive (SRA) under BioProject under accession PRJNA587717. Sliding window files of aforementioned sizes and unmappable read files are available through the corresponding author.

