## Supplementary material for "Netome: The Molecular Characterization of Neutrophil Extracellular Traps (NETs)": u_46267_0_supp_1031906_q7mzdz_version2

**Running Title:** Genomics and Proteomics of NETs

**Authors:**

David Scieszka[1], Yi-Han Lin[2], Weizhong Li[1], Saibyasachi Choudhury[1], Yanbao Yu[2], Marcelo Freire[1,2,3*]

**Affiliations:**

[1]Department of Human Biology and Genomic Medicine, J. Craig Venter Institute, La Jolla, CA 92037, USA.

[2] Department of Infectious Disease, J. Craig Venter Institute, Rockville, MD 20850, USA.

[3] Department of Infectious Diseases, University of California San Diego, La Jolla, California, USA.

* Corresponding author
Marcelo Freire, D.D.S., Ph.D., D.Med.Sc
Associate Professor
Genomic Medicine and Infectious Diseases
4120, Capricorn Lane, 92037
La Jolla, CA


Supplementary Material

**Supplementary Figure 1**

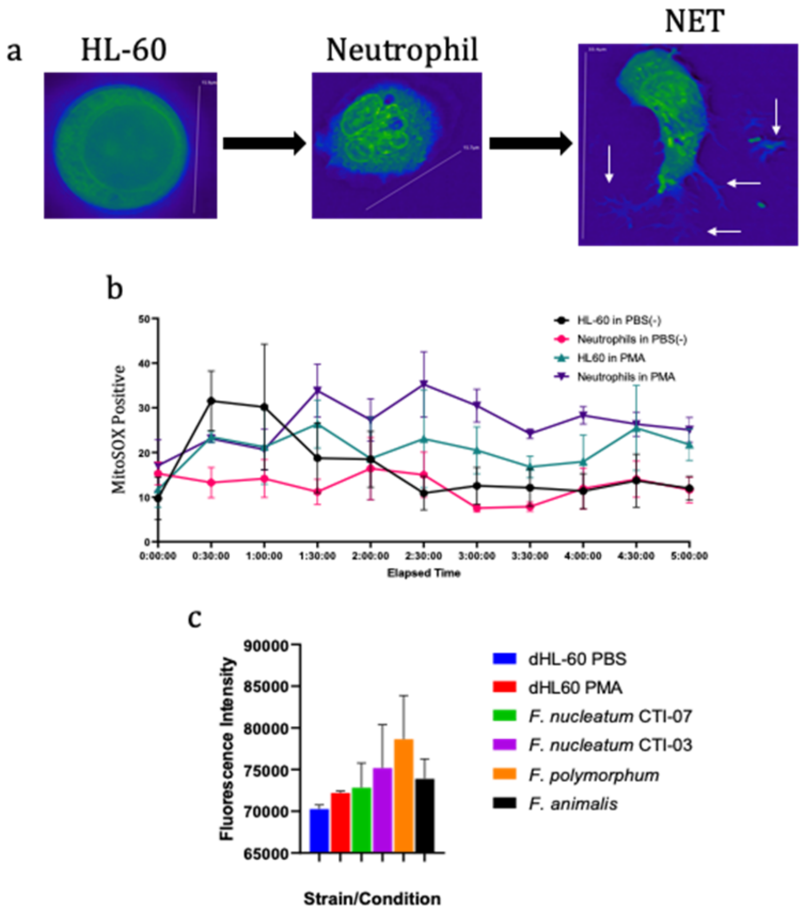


**Supplementary Figure 1. HL-60 differentiation, NET formation, and coculture. (a)** Holotomographic microscopy images, digitally stained based on RI (refractive index) confirms the differentiation of HL-60 cells to neutrophils (dHL-60) after incubation with MA, DMSO, LPS, or formaldehyde, and successful release of NETs after coincubation with different strains of bacteria. Arrows point to neutrophil extracellular DNA or bacterial DNA biolayer **(b)** Quantification of flow cytometry utilizing MitoSOX stain under different conditions of PMA or PBS. Maximum neutrophil ROS generation is seen at 2.5 hours (n=3). PMA final concentration 1,000 nM. **(c)** Quantification of neutrophil elastase through fluorescent activity immediately after coculture with gram negative *Fusobacterium* strains: *polymorphum, animalis,* and *nucleatum* substrains CTI-03 and CTI-07.

**Supplementary Figure 2**

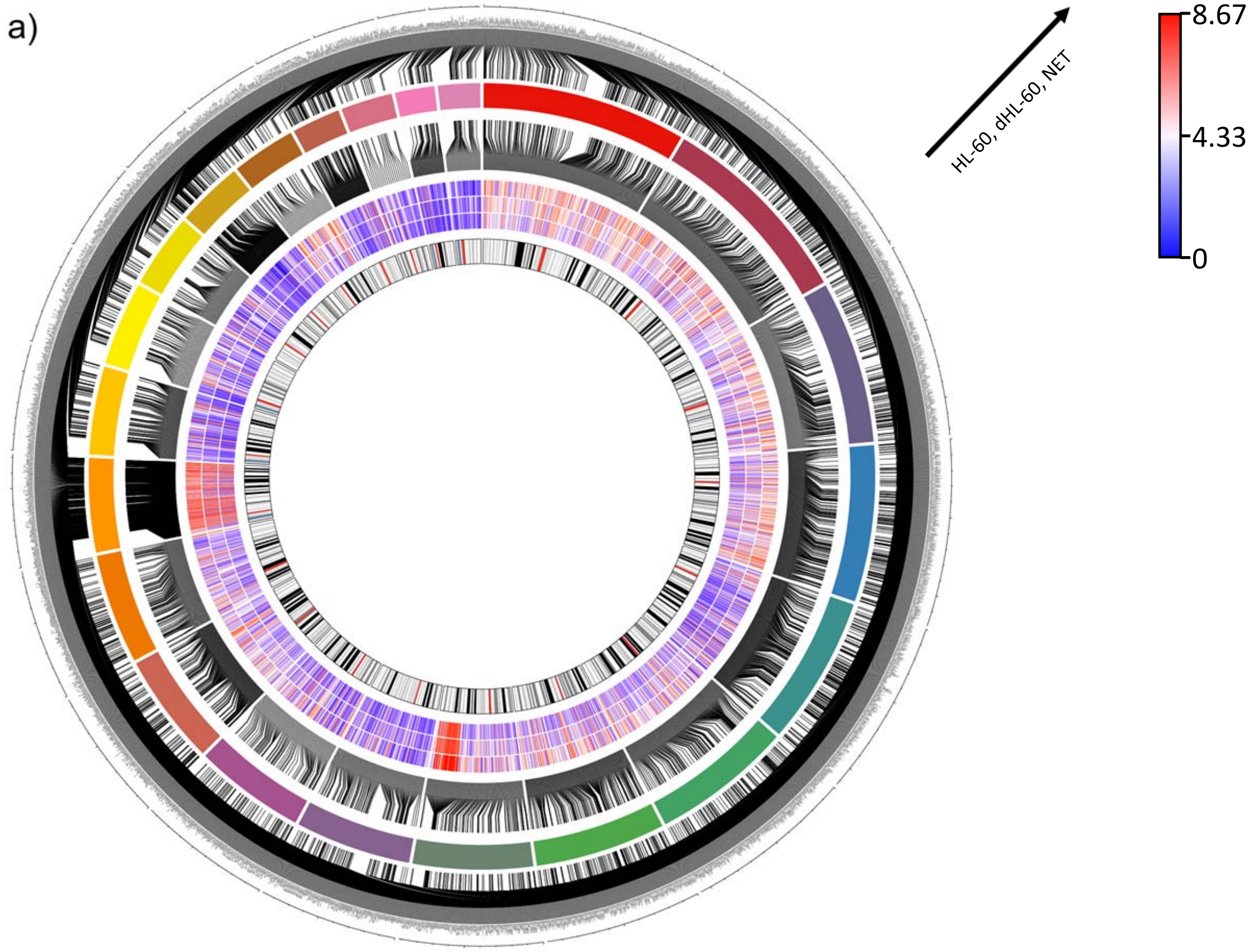


**Supplementary Figure 2. Circos plot of genome-wide scan.** Genes shown were generated from the 25,488 regions that exhibited consistent 1.5-fold enrichment or depletion in the NET sample relative to HL-60 and dHL60. Enrichment/depletion levels were binned into 26 quantiles ranging from 0 to 8.66 for aesthetic reasons.

**Supplementary Figure 3**

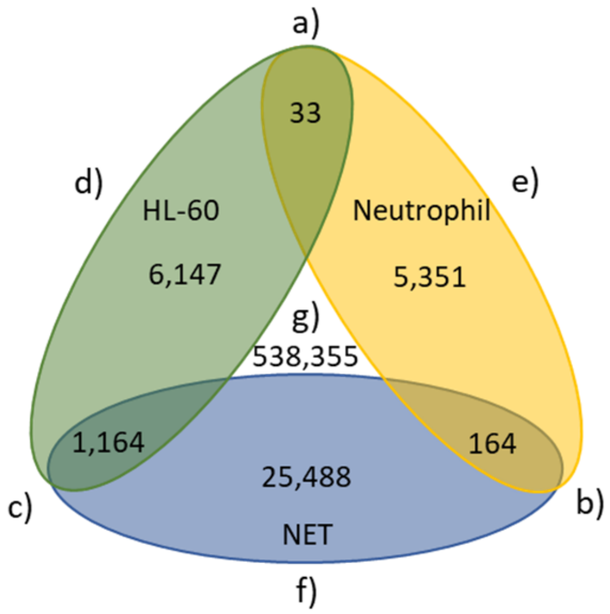


**Supplementary Figure 3. Comparisons using the 5,000nt sliding window between individual samples and averaged sample enrichments. (a-c)** Averages of the two adjacent samples were compared. **(d-f)** Single sample compared to the unaveraged grouping of the other two samples. **(a)** Average of HL-60 and neutrophil samples showed enrichment/depletion in 33 regions over NET sample. **(b)** Average of NET and neutrophil samples showed enrichment/depletion in 164 regions over HL-60 sample. **(c)** Average of NET and HL-60 samples showed enrichment/depletion in 1,164 regions over neutrophil sample. **(d)** HL-60 sample showed consistent enrichment/depletion in 6,147 regions when compared to the grouping of neutrophil and NET. **(e)** Neutrophil sample showed consistent enrichment/depletion in 5,351 regions when compared to the grouping of HL-60 and NET. **(f)** NETs are enriched/depleted in 25,488 regions when compared to the grouping of HL-60 and neutrophil. **(g)** There are 538,355 regions that show no consistent enrichment or depletion among any sample when compared to the grouping of the other two.

**Supplementary Figure 4**

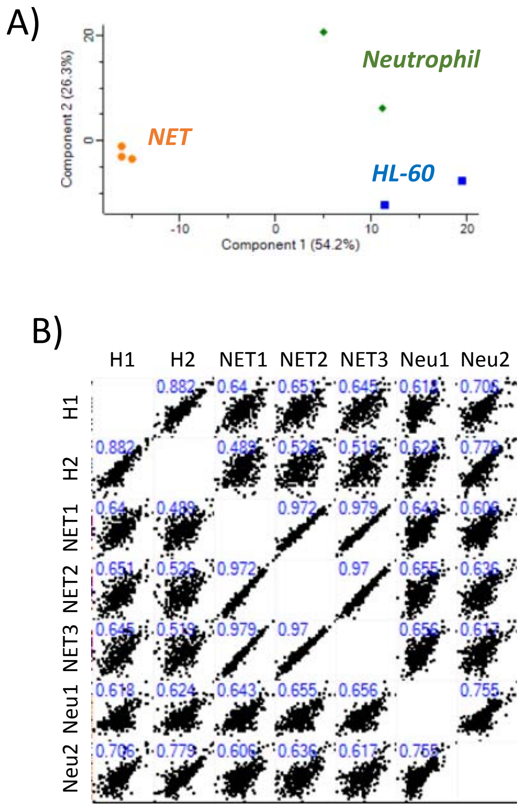


**Supplementary Figure 4. Comparison of NET to differentiated and undifferentiated cells. (A)** proteomic principal component analysis demonstrating clusters of NET groups in comparison to differentiated neutrophils and undifferentiated cells (HL-60). **(B)** Pearson correlation values were displayed to each analysis.

**Supplementary Figure 5**

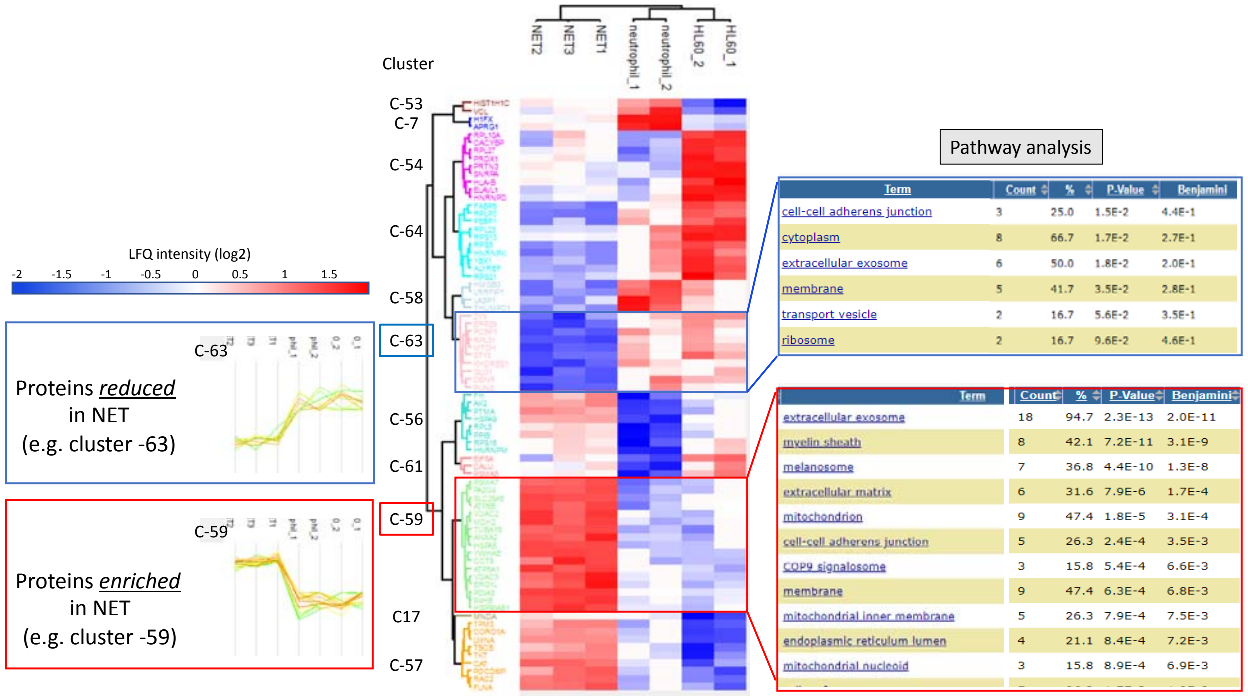


**Supplementary Figure 5. Comparison of enriched proteins in NETS.** In comparison to neutrophils and HL60 cells, 75 proteins were enriched (downregulation in blue, upregulation in red). Pathway analysis demonstrated top functions involved in proteins enriched in NETs.

**Supplementary Figure 6**

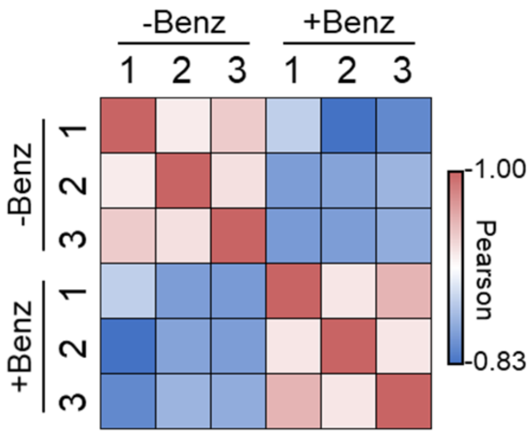


**Supplementary Figure 6.** Correlation analysis of the six NET samples (n = 3 for each group, with or without benzonase treatment). Pearson correlation values were color coded as indicated in the scale bar.
